# The Evolution of Microbiome-Mediated Traits

**DOI:** 10.1101/2024.03.29.587374

**Authors:** Bob Week, Andrew H. Morris, Brendan J. M. Bohannan

## Abstract

The role of host-associated microbiomes in shaping host fitness is becoming increasingly recognized, and may have important evolutionary consequences. The degree to which microbiomes contribute to host evolution depends on how they are inherited. To understand the conditions under which host evolution is mediated by microbiome inheritance, we introduce a simple mathematical model. Our model tracks the evolution of a quantitative host trait that is mediated by host genotype and host microbiome. Unlike host genotype, the host microbiome may have several routes of transmission, including social contact and environmental acquisition. Our model takes a breeding-value approach to formally generalize classical quantitative genetic theory to account for these various modes of microbiome inheritance. A key prediction of our model is that, as long as microbiomes are transmitted between host generations, their mode of transmission has no effect on host trait dynamics. To understand the conditions required for this result, we analyze microbiome-explicit simulations that break important model assumptions. We then discuss avenues to generalize our model and connections with related concepts. Taken together, this work provides novel insights into the dynamics of non-Mendelian populations, and makes first steps towards establishing a theory of microbiome-mediated trait evolution.

## Introduction

There is a rapidly growing body of evidence demonstrating that microbiomes mediate many important host traits (Friesen et al. 2011; McFall-Ngai et al. 2013). This realization, combined with recent recognition that host-associated microbiomes can be transmitted across generations, has spurred interest in understanding how microbes may determine host trait dynamics (Russell 2019; Leftwich et al. 2020; Murphy et al. 2023). Approaches to study this emerging topic draw on insights from the intersection of evolutionary and microbiome biology, and have important applications in human health (Gilbert et al. 2018), agriculture (Gopal and Gupta 2016), and conservation (Trevelline et al. 2019; Mueller et al. 2020). Progress in understanding the dynamics of host-microbiomes has been made recently using simple mathematical models (Vliet and Doebeli 2019; Leftwich et al. 2020; Bruijning et al. 2021). However, these studies have focused primarily on understanding microbiome dynamics, rather than the consequences of microbiome dynamics for microbiome-mediated host traits. Hence, there remains a gap in modeling approaches for predicting host trait dynamics in response to microbiome dynamics. To fill this gap, we introduce a quantitative genetic model of host traits that accounts for the effects of dynamic host microbiomes.

Our approach leverages insights gained from several recent theoretical advances in the study of host-microbiome evolution. For example, Zeng and Rodrigo (2018) took a metacommunity approach to simulate the dynamics of microbiomes comprised of several microbe species within a host generation. They found that parent-offspring vertical transmission leads to increased variation of microbiomes among hosts, and that environmental acquisition of microbes leads to decreased variation among hosts. In another study, van Vliet and Doebeli (2019) investigated a multi-level selection model tracking the relative abundances of two microbial strains. The authors found that microbes may evolve to benefit their hosts when host generation time is short relative to the timescale of microbial evolution, and when microbes are passed vertically from parent to offspring. Leftwich et al. (2020) studied two models tracking the presence of a single microbe species in a host population, and concluded that the efficiency of vertical transmission of microbes plays a more important role than host selection for establishing host-microbe associations.

More recently, Roughgarden (2023) introduced a mathematical framework to model host-microbiome integration that makes comparisons with population genetic theory. A particularly important contribution made by Roughgarden’s paper, which we incorporate into our model, is a generalization of the concept of inheritance. Nuclear genes are transmitted vertically from parent to offspring, leading to what Roughgarden (2023) calls “lineal inheritance”. On the other hand, microbiomes can be horizontally transmitted through social contact or environmental acquisition (Raulo et al. 2021; Tavalire et al. 2021), and can lead to a resemblance of parent and offspring populations (but not necessarily individuals) that forms what Roughgarden (2023) calls “collective inheritance”. We adopt this terminology in this paper. As Roughgarden (2023) points out, systems exhibiting collective inheritance are capable of evolutionary descent through modification, and may therefore respond to selection.

The approaches to modeling host-microbiome evolution mentioned above focus directly on the dynamics of microbiome composition. However, the consequences of microbiome change for microbiome-mediated host traits are not obvious. To establish this connection, a quantitative genetic approach may be useful, as several authors have recently suggested (Bruijning et al. 2021; Henry et al. 2021; Mueller and Linksvayer 2022). Henry et al. (2021) proposed that the classical decomposition of host phenotypic variance into genetic and environmental components can be extended to include a component for the host microbiome. This decomposition allows for the formal definition of a microbiome-mediated trait as a quantitative trait of the host with variance explained in part by variation of host-associated microbiomes. A simulation model of microbiome-mediated trait dynamics was studied by Bruijning et al. (2021). There, the authors found that fluctuating selection decreases vertical transmission of microbiomes. Following these studies, Mueller and Linksvayer (2022) further developed a conceptual foundation for understanding the eco-evolutionary processes involved in microbiome-mediated trait dynamics and their use for microbiome manipulation.

Building on these previous quantitative genetic perspectives, we introduce a mathematical model of microbiome-mediated trait dynamics. Our model accounts for vertical transmission of microbiomes from parent to offspring (corresponding to Roughgarden’s “lineal inheritance”), shedding of microbes into the host environment, and environmental acquisition. Collective inheritance occurs when host offspring acquire microbes from the environment that ultimately descend from a parental microbiome. With this model of microbiome transmission, we track the inheritance of additive genetic and additive microbial effects on host traits. Compared with previous approaches to model host-microbiome evolution, this approach is unique because it allows us to apply standard quantitative genetic assumptions to model the response of a microbiome-mediated trait to selection.

Our paper begins by introducing our analytical model. To test predictions made by our analytical model when assumptions are broken, we introduce our approach to simulate a microbiome-explicit model. We then describe our approach to compare the two models across sets of randomly drawn model parameters and present our results. Finally, we discuss limitations and possible generalizations of our analytical approach, and compare the concept of a microbiome-mediated trait with phenotypic plasticity and niche construction.

## Methods

### Analytical Model

Our analytical model considers an infinite population of asexually reproducing host organisms that have discrete, non-overlapping generations. The life-cycle of the host begins at birth with instantaneous colonization by microbes. As the host matures, host microbiomes develop independently. Host traits are determined by their microbiomes at adulthood. Viability selection on host traits occurs after hosts have matured. After selection, adult hosts replace themselves with offspring and shed their microbiomes into the environment, at which point the life-cycle repeats. An overview of our model is illustrated in Figure 1.

**Figure 1:**
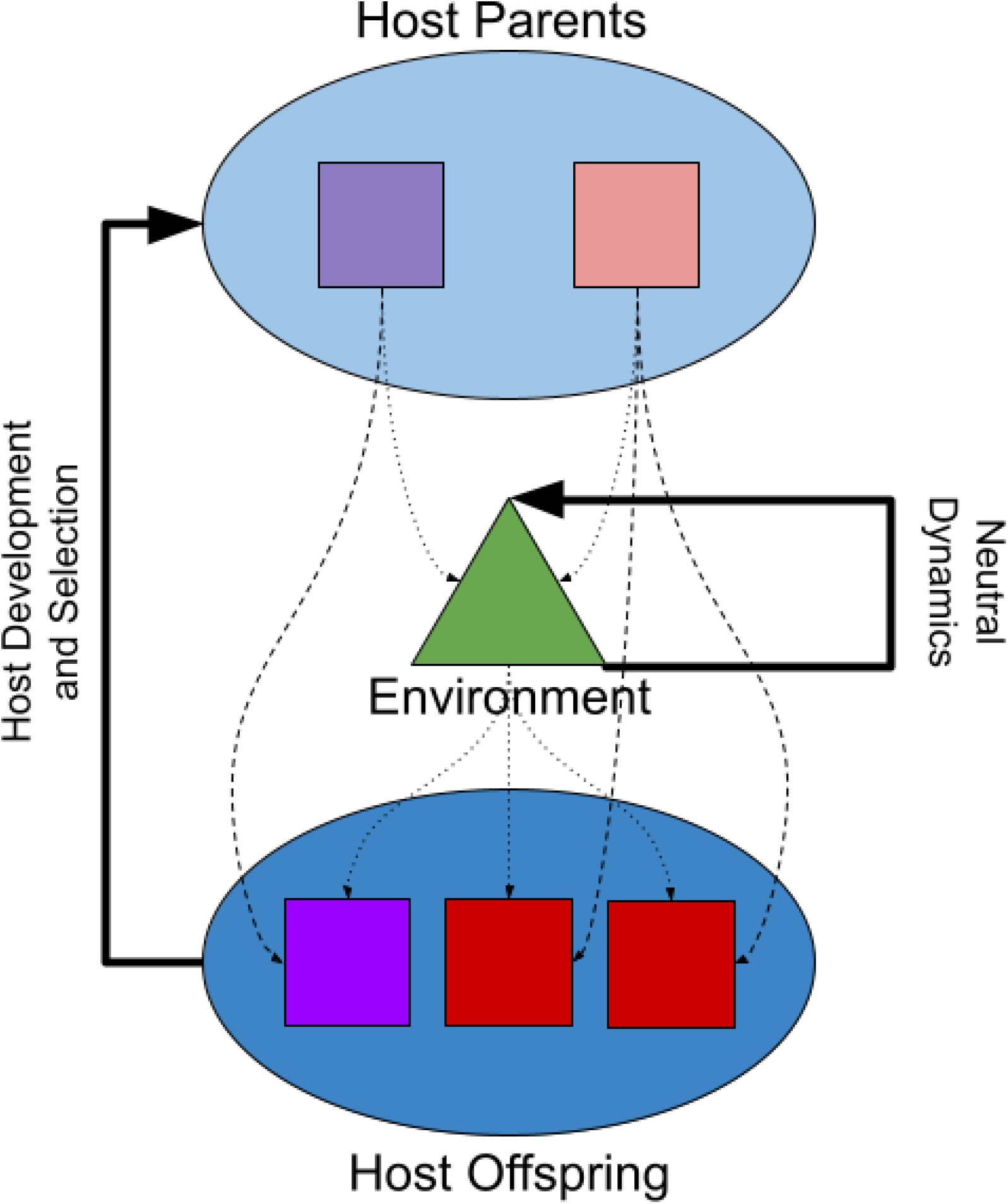
An illustration of the general structure of our model over a single host generation. Hosts are represented by squares, with trait value encoded by color. The light blue oval at the top of the digram contains the host parents (the light purple and light red squares). The dark blue oval at the bottom of the diagram contains host offspring (the dark purple and two dark red sqaures). The green triangle in the center of the digriam represents the environment into which the host parents shed microbes, and from which host offspring acquire their microbes. The dashed lines indicate host ancestry, and form lines of lineal inheritance of microbes and genes. The dotted lines indicate environmentally mediated transmission of microbes, which form lines of collective inheritance. The bold arrow to the left indicates components of the host life-cycle (development and selection) not depicted in this diagram. Selection is based on viability, which explains why there are fewer host parents than host offspring. The bold arrow to the right indicates that the environmental microbiome undergoes neutral stochastic dynamics while the hosts undergo development and selection.

We have strived to create the simplest mathematical representation of microbiome transmission, establishment and maturation, and subsequent selection on microbially-mediated traits, Our base model can be used as a foundation for more complex models, for example by adding sexually-reproducing hosts. However, because we focus on the evolution of host mean trait rather than host trait variance, a model of random mating will likely yield identical results to our simpler asexual model.

### Host Trait Architecture

Host traits are assumed to decompose into three additive components: *z* = *g* + *m* + *e*, with *z* being the trait value for an individual host organism. The first component, *g*, is the sum of additive effects across all genetic loci of the host, which is commonly known as a “breeding value” (Lynch and Walsh 1998; Walsh and Lynch 2018). However, for the sake of clarity, we adopt a less common convention and refer to *g* as a “additive genetic value” (Roughgarden 1979; Bulmer 1980). The second component, *m*, is the sum of additive effects across all genetic loci of microbes that comprise the host microbiome, which we call the additive microbial value in analogy to *g*. The third component, *e*, is an unbiased error term that accounts for additional trait variation, such as microenvironmental variation and random developmental noise (Bulmer 1980; Lynch and Walsh 1998). We further assume the components *g, m* and *e* are independent and normally distributed among host individuals with respective population means 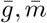, 0 and variances *G, M, E*. Host population trait mean and variance are then given by 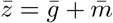 and *P* = *G* + *M* + *E*. Combining this notation with our definition of a microbiome-mediated trait, the host trait is a microbiome-mediated trait whenever *M >* 0.

### Selection

We assume viability selection acts on host traits and write the fitness of a host with trait value *z* as *W* (*z*). The post-selection mean trait of the host population, 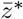, is given by the size biased average

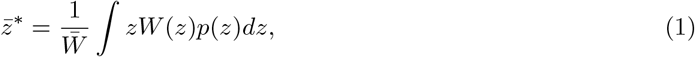

where *p*(*z*) is the distribution of trait values in the host population and 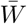 is host mean fitness. When traits are normally distributed, and selection is frequency-independent (Lande 1976), the difference between post-selection and pre-selection mean traits (called the selection differential) can be expressed as

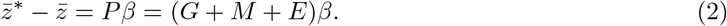

Here, *β* (called the selection gradient) is the derivative of the log of mean fitness with respect to mean trait (Lande and Arnold 1983), which indicates how mean fitness changes as we vary mean trait. The consequences of selection for host trait variance can also be calculated, and generally lead to non-random associations between additive genetic and microbial values. However, for simplicity we focus our analytical model on the dynamics of host mean trait.

Equation (2) states that selection acts to shift the mean trait of a population in a direction that maximizes fitness (accounted for by the selection gradient) by a magnitude proportional to the phenotypic variance of that population. However, Equation (2) does not describe the hosts evolutionary response to selection because the components of phenotypic variance are not all perfectly inherited. To determine the degree to which each component of trait variation is inherited by host offspring, we need to model the transmission of each factor.

### Routes of Transmission

Genetic factors are lineally inherited, being transmitted strictly from parent to offspring. We assume that trait variation due to environmental noise is not inherited so that the value of noise in an offspring is statistically independent from the value of noise in that offspring’s parent. Microbiome transmission is more complicated, possibly following several different routes simultaneously. Here, we describe our approach to taming the complexity of microbiome transmission while also retaining its most salient features so that our framework can provide a mathematically tractable, yet non-trivial description of microbiome-mediated trait dynamics.

We make the simplifying assumption that hosts are instantaneously colonized by microbes at birth. Additionally, we assume that a microbe inhabiting a host was inherited from that hosts immediate parent with probability *𝓁*, and acquired from the host environment with probability 1 − *𝓁*. The environment is given a hypothetical additive microbial value, *ξ*, which is the additive microbial value of an adult host when the composition of that hosts microbiome is equivalent to that of the environment.

Following our assumptions, the expected trait of a host offspring born to a parent with additive genetic value *g* and additive microbial value *m*, and into an environment with additive microbial value *ξ*^*′*^, is given by

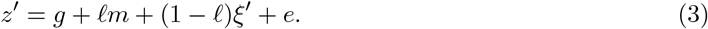

For simplicity, we assume the absence of fecundity selection so that each host that survives viability selection has the same expected number of offspring. Then averaging across the offspring population yields

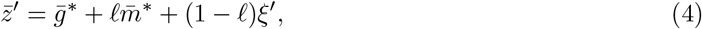

where 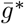 and 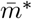 are the mean additive genetic and microbial values in the post-selected parental host population.

We account for the possibility that microbes acquired from the environment by host offspring are ultimately derived from host parent microbiomes. That is, we allow for the environment experienced by host offspring to be shaped by host parents via microbial shedding. To model this, we assume microbes in the environment experienced by host offspring either come directly from the environment experienced by host parents, or from the pooled microbiome of selected host parents. Figure 1 presents a path diagram to visualize the different modes of microbiome transmission in our model.

We denote by *ε* the proportion of the microbiome in the environment experienced by host offspring that come directly from the environment experienced by host parents. We assume a closed environment so that 1 − *ε* is the proportion of the environmental microbiome that comes from the pooled microbiome of selected host parents. The mean trait of the host offspring population is then given by

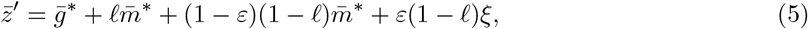

where *ξ* is the additive microbial value of the parental host environment. Before we compute the change in host mean trait across successive host generations, we first discuss genetic mutation and microbiome dynamics.

### Mutation and Microbiome Dynamics

In our framework, focused on host trait evolution, within-host microbiome dynamics acts as the microbiome analog of genetic mutation. While mutation results in random differences between parent and offspring genotypes, we assume microbiome dynamics results in random differences between host microbiomes at birth and at adulthood. This conceptual similarity has also been pointed out by Bruijning et al. (2021). Both genetic mutation and microbiome dynamics can contribute to variation of microbiome-mediated traits. In the following two paragraphs, we provide conceptual snapshots for how these processes are treated in our framework.

### Mutation

In classical quantitative genetics, mutation has been modeled by assuming offspring additive genetic value *g*^*′*^ is a random variable that may depend on parental additive genetic value *g* (Lande 1975; Turelli 1984, 2017; Bürger 2000; Barton et al. 2017). Our analytical model assumes the effects of mutation are unbiased so that offspring additive genetic value equals parental additive genetic value in expectation; *E*[*g*^*′*^] = *g*. Combining this assumption with our assumption that the host population is infinitely large, the mean additive genetic value of the host offspring population, 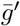, is a deterministic quantity equal to the mean additive genetic value of the post-selected host parental population, 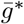. In this case, mutation has no effect on mean trait evolution between consecutive host generations.

### Microbiome Dynamics

The effect of microbiome dynamics on additive microbial values is modeled in analogy to the effect of mutation on additive genetic values, but with several important differences. For instance, instead of comparing additive values between parent and offspring, we compare microbial additive values before and after host development. In particular, we introduce the concept of an initial host additive microbial value, *m*_0_, which is the additive microbial value of a hosts microbiome immediately after its assembly during host birth. Because, in our model, traits are only expressed by mature hosts, *m*_0_ is a hypothetical value, equal to the additive microbial value of an adult host that has an identical microbiome composition of the newborn host. Similar to our assumption on the effects of genetic mutation, we assume the additive microbial value of an adult host, *m*, is equal to its initial additive microbial value on average; *E*[*m*] = *m*_0_. This assumption precludes the possibility for microbiome dynamics to bias host mean trait dynamics.

In addition to within-host microbiome dynamics, our model tracks the additive microbial value of the environment, which has no genetic analog. The environmental microbial value of the host population before development is denoted by *ξ*_0_, and the analogous value for the host population after development is denoted by *ξ*. We assume *ξ* is a random variable with a mean equal to *ξ*_0_. This creates a source of environmental stochasticity that can result in random fluctuations of host mean trait value even when the host population is assumed to be infinitely large, and is closely related to stochastic selection (Gillespie 1972; Felsenstein 1976; Lande 2007; Chevin 2019; Cotto and Chevin 2020). An important difference is that, while stochastic selection results from environmental stochasticity affecting fitness as a function of trait value, here environmental stochasticity affects trait value as a function of the environment (considering host-associated microbiomes as components of the host environment). However, if we were to consider fitness solely a function of additive genetic value, the effects of environmental stochasticity caused by environmental microbiome dynamics would result in stochastic selection.

Combining our treatment of mutation and microbiome dynamics with our model of host microbiome assembly (illustrated by the directed graph in Figure 1), the expectation of a host offspring’s trait value at maturity, conditioned on knowing the environmental additive microbial value in the parental generation, *ξ*, is given by

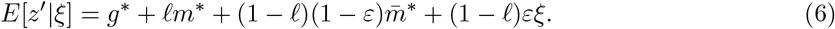

Averaging across matured offspring provides the mean trait in the offspring generation,

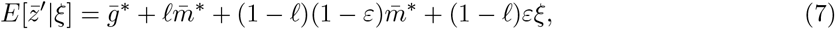

which is in agreement with Equation (5) obtained above. Hence, our approach to modeling microbiome dynamics does not alter mean trait dynamics in expectation. However, environmental microbiome dynamics creates uncertainty in the offspring mean trait value. More precisely, using the law of total expectation, the expression for 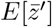 matches that for 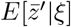, except for replacing *ξ* with *ξ*_0_, the initial environmental additive microbial value in the parental generation. Then, using the law of total variance, the variation in offspring mean trait due to environmental microbiome dynamics is given by

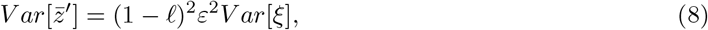

where *V ar*[*ξ*] is the variance of the environmental additive microbial value due to environmental microbiome dynamics. We show how to calculate this variance using details from our simulation model below in the subsection titled *Microbial Architecture of Host Trait*.

### Mean Trait Dynamics

Now that we have accounted for trait architecture, selection, transmission, mutation, and microbiome dynamics, we are ready to calculate the change in host mean trait between successive generations. The difference between offspring and parental population mean trait values is denoted by 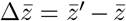. In this subsection we ignore the stochastic effects of environmental microbiome dynamics by equating 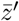 with 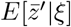. Setting *κ* = (1 − *𝓁*)(1 − *ε*) and applying Equation (5), we find

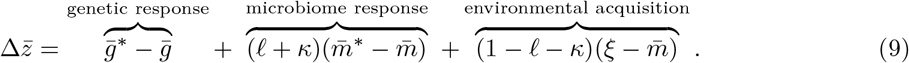

Equation (9) demonstrates that the change in mean trait between successive generations can be partitioned into three components. The first component is the genetic response to selection, the second component is the microbiome response to selection, and the third component is due to environmental acquisition. Connecting our model with the notions of lineal and collective inheritance introduced by Roughgarden (2023), Equation (9) also shows that the response to microbiome selection is mediated by lineal inheritance, captured by parameter *𝓁*, and collective inheritance, captured here by parameter *κ*. Together, 0 ≤ *κ* + *𝓁* ≤ 1 determine the proportion of additive microbial variation in the parental population that is inherited by the offspring population.

The expression for mean trait dynamics can be simplified using our assumption that additive genetic and microbial values are normally distributed. This allows us to write differences between pre- and post-selected mean additive values as products of the selection gradient *β* and variances of additive values, similar to Equation (2). In particular, combining lineal and collective inheritance into a single parameter *v* = *𝓁* + *κ*, Equation (9) can be rewritten as

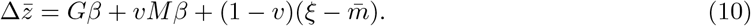

Equation (10) generalizes the classical quantitative genetic expression 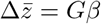, the univariate case of Equation (7a) in Lande (1979). These expressions coincide when two conditions are satisfied: 1) when *v* = 0, so that *M* is not heritable, and 2) when 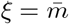, so that the composition of the environmental microbiome matches the average composition of the host microbiome.

In the case that all of *M* is heritable, so that *v* = 1, Equation (10) simplifies to

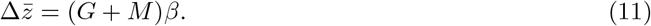

This occurs when all of the microbes acquired by host offspring ultimately derive from the pooled microbiome of the selected host parents. This can happen in several possible ways. For instance, it’s possible that host offspring acquire their entire microbiome from their immediate parent so that microbiomes are strictly lineally inherited, corresponding to *𝓁* = 1 and *κ* = 0. At the opposite extreme, this occurs when microbiomes are strictly collectively inherited, corresponding to *𝓁* = 0 and *κ* = 1. More generally, Equation (11) holds whenever microbiome acquisition is some combination of lineal and collective inheritance, corresponding to *𝓁* + *κ* = 1. Importantly, this indicates that the response to selection does not depend on whether inheritance is lineal or collective.

Even when the host microbiome is not at all heritable, so that *v* = *𝓁* + *κ* = 0, our model implies host trait dynamics can still be mediated by microbiomes. In this case, Equation (10) simplifies to

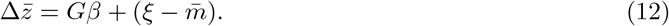

Here, the host offspring acquire their entire microbiome from their environment so that 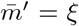. If the microbiome of the environment experienced by host offspring is compositionally distinct from the pooled microbiome of selected host parents, then 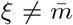 and the host mean trait exhibits a plastic response to environmentally acquired microbes. We elaborate on this connection with phenotypic plasticity and reaction norms in the discussion section below.

To complete the description of our model, we also track the expected dynamics of the environmental microbial value, which are given by

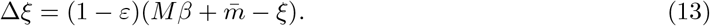

This equation states that the environmental microbial value moves towards the post-selection additive microbial value 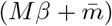 at rate determined by host shedding. In particular, when parental hosts do not shed microbes into the offspring environment (i.e., when 1 − *ε* = 0), there is no expected change in the environmental microbial value.

Our analytical model is sufficiently tractable to provide several novel insights into the dynamics of microbiome-mediated traits, but relies on a series of important simplifying assumptions. Two major assumptions involve the microbial architecture of host traits and infinite host population size. In the following section, we introduce microbiome-explicit simulations to investigate the consequences of breaking both of these assumptions.

### Microbiome-Explicit Simulations

To investigate the robustness of our analytical predictions, we simulated a microbiome-explicit model. The simulation model builds directly on our analytical model by incorporating additional details that allow us to break two important assumptions. In particular, we explicitly track abundance dynamics of microbial species in each microbiome. Each microbe species is associated with a per capita additive effect on host trait, which we use to compute the additive microbial values. These details create the microbial architecture of the host trait. We also assume a finite host population size, which results in random genetic drift of host genotypes and, in analogy, random drift of host-associated microbiomes. Below we describe the details for the microbial architecture of host traits and dynamics of finite host populations implemented in our simulation model. We then compare simulated results with analytical predictions.

### Microbial Architecture of Host Trait

We take Hubbell’s classical approach to model microbiomes as dynamic ecological communities (Hubbell 2011). This is the simplest approach to modeling microbiome dynamics; in addition, neutral models have been shown to capture important aspects of microbiome dynamics in a number of host species (Burns et al. 2015; Venkataraman et al. 2015; Heys et al. 2020; Henry and Ayroles 2022). This implies a fixed number *J*_*e*_ of microbe individuals in each microbiome, and stochastic fluctuations in species’ abundances without bias for any particular species. The *J*_*e*_ microbe individuals are assumed to be subdivided into *S* microbial species. The abundance of microbe species *i* is denoted *n*_*i*_ so that 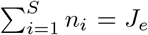. In our model, a microbiome is completely described by the abundances of microbial species that inhabit it, and we write (*n*_1_(*τ*), …, *n*_*S*_(*τ*)) as the vector of abundances at host age *τ* . The neutral community dynamics are modeled by assuming the microbiome at time *τ* + 1 follows a multinomial distribution with *J*_*e*_ trials, *S* mutually exclusive events, and probability of event *i* equal to *n*_*i*_(*τ*)*/J*_*e*_. This process iterates *L* times during host development so that *τ* = 0, 1, …, *L*. Initial microbiomes are determined by the assembly process described above for our analytical model. In particular, each of the *J*_*e*_ microbes chooses its ancestry following the path diagram shown in Figure 1. The parameter *J*_*e*_ plays the analogous role as effective population size in models of population genetics by determining the rate of stochastic dynamics (Wakely 2016).

We associate each microbial species with a per capita additive effect on host trait. The per capita additive effect for microbial species *i* is denoted by *α*_*i*_. The additive microbial value of an adult host, with microbiome (*n*_1_(*L*), …, *n*_*S*_(*L*)), is then given by

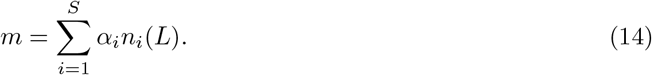

Hubbell’s model of neutral community dynamics implies *E*[*n*_*i*_(*L*)] = *n*_*i*_(0), so there is no bias in the change in microbial species abundances. Combined with our model of host trait microbial architecture, this implies that our simulation model of microbiome dynamics does not bias host trait evolution. More precisely, setting *m*_0_ as the additive microbial value of the initial host microbiome (*n*_1_(0), …, *n*_*S*_(0)), we have *E*[*m*] = *m*_0_.

To account for the stochastic dynamics of microbial communities while comparing simulated results with our analytical expectations, it is useful to know the variance of microbiome dynamics. That is, the variance of the difference of additive microbial values before and after host development; *V ar*[*m* − *m*_0_]. Using the details provided above, this variance is given by

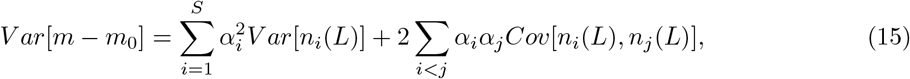

where

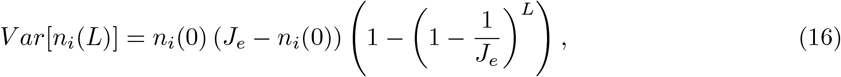

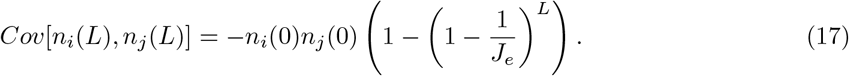

Because the model of neutral community dynamics we adopt is mathematically equivalent to the multiallelic Wright-Fisher model (Ewens 2004), many insights carry over. For example, we see that, as *J*_*e*_ increases (playing an analogous role as effective population size), the variance of microbiome dynamics decreases. However, this model of microbiome dynamics has a novel consequence for host trait variation. Our analytical model assumed the effect of microbiome dynamics on host trait to be identical for each host individual. In contrast, the description of neutral community dynamics above, Equations (16) and (17) in particular, show the random effects of microbiome dynamics depend on the composition of the hosts initial microbiome.

### Finite Host Population Size

In addition to stochastic microbiome dynamics resulting from finite community size, we also assume a finite number of host individuals, which leads to another source of stochasticity we call host drift. Host drift encapsulates both classical random genetic drift of host allele frequencies, and an analogous process for host-associated microbiomes. As a direct generalization of random genetic drift, host drift reduces the efficacy of selection and erodes heritable genetic variation (Bulmer 1980; Barton and Turelli 2004; Walsh and Lynch 2018). By the same mechanism, host drift also erodes heritable phenotypic variation explained by host-associated microbiomes.

Our approach to modeling finite host population size is similar to the Wright-Fisher model with selection (Etheridge 2011). In particular, we assume the host population consists of *N*_*e*_ individuals in each generation. Each of the *N*_*e*_ host offspring in the following generation pick their parents independently at random, with probability proportional to the fitness of each parent. That is, if the *i*th individual in the parental population has trait value *z*_*i*_, and therefore fitness *W* (*z*_*i*_), an offspring will choose this parent with probability

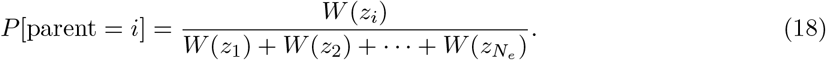

During host reproduction, we assume the selected parents shed microbes into the environment experienced by newborn host offspring. The formation of the environmental microbiome that host offspring are exposed to follows the same process outlined for our analytical model in the subsection titled *Routes of Transmission*. In particular, the pooled parental microbiome consists of the microbiomes of individuals in the parental population selected as parents by host offspring. This implies each of the *J*_*e*_ microbes in the offspring environment pick their ancestry at random, with probability *ε* descending from the parental host environment, and probability 1 − *ε* descending from the pooled microbiome of selected host parents. Initial offspring microbiomes are then formed, also following the process outlined by our analytical model, where each of the *J*_*e*_ microbes in an offspring microbiome is acquired from that offspring’s immediate parent with probability *𝓁*, and from the environment with probability 1 − *𝓁*. Unlike our analytical model, however, in our simulation model we also account for the species identity of each microbe. The species of a microbe is determined by the composition of the microbiome it originated from. In particular, following our notation from the subsection titled *Microbial Architecture of Host Trait* above, if the microbe originated from a microbiome described by the vector of abundances (*n*_1_, …, *n*_*S*_), then it belongs to species *i* with probability *n*_*i*_*/J*_*e*_.

To complete the specification of our approach to modeling finite host population size, we make additional assumptions on the mechanics of host genetic mutation. While our analytical model only assumes the effect of mutation is unbiased, without assuming any particular distribution, our simulation model follows the Gaussian allelic approximation, which implies the effects of mutation on additive genetic values are normally distributed (Kimura 1965; Lande 1975; Bürger 2000). We denote the variance of mutation by *µ*. Because our simulation model assumes finite host population size, mutation provides an additional source of stochasticity for host trait evolution. In particular, denoting 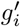 the additive genetic value of offspring *i* and *g*_*i*_ the additive genetic value of that offspring’s parent, our assumptions imply the effect of mutation on host mean trait, given by 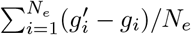, has expectation zero and variance *µ/N*_*e*_. Mutation does not appear as a source of stochasticity in our analytical model, because there we assume infinite host population size. To illustrate our simulation model, a set of 100 replicated runs across 200 host generations is presented in Figure 3.

**Figure 2:**
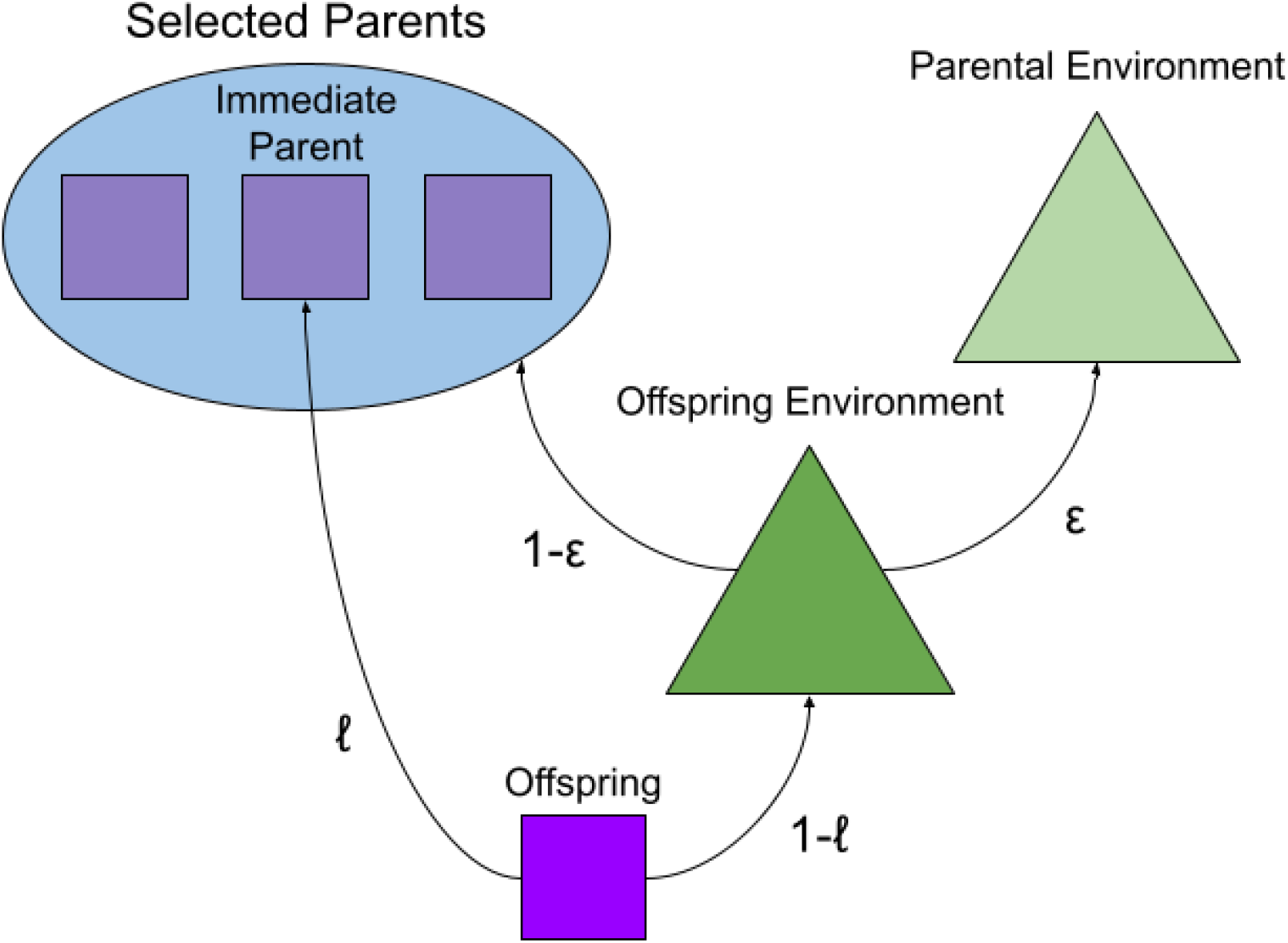
Different modes of microbiome inheritance. The arrows indicate different routes of microbial ancestry, and are labelled by their probability. A microbe in the offspring microbiome descends from the microbiome of that offsprings immediate parent with probability *𝓁*, and from the offspring environmental microbiome with probability 1 − *𝓁*. In turn, a microbe in the offspring environmental microbiome descends from the parental environmental microbiome with probability *ε*, and from the pooled parental microbiome with probability 1 − *ε*.

**Figure 3:**
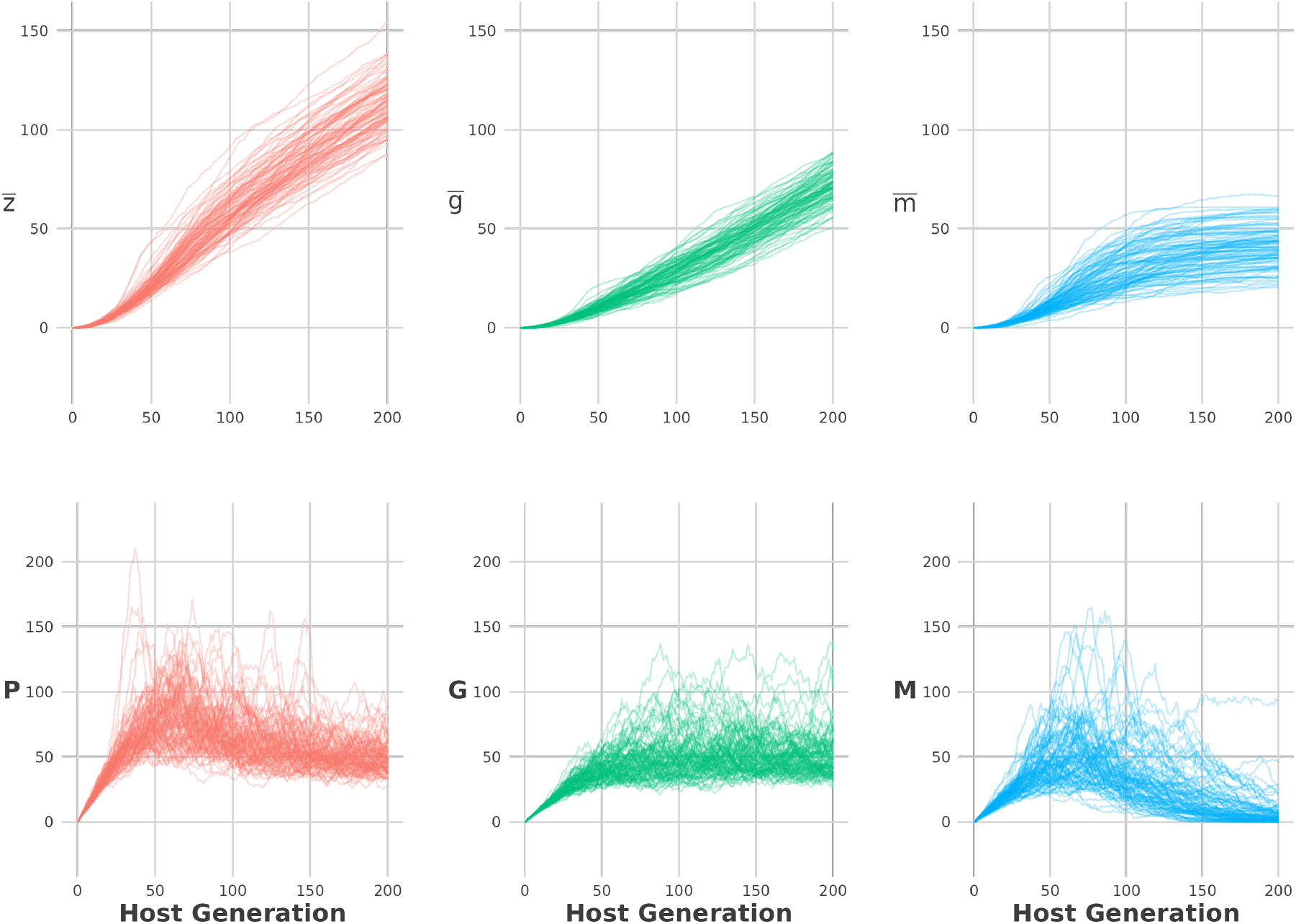
A set of 100 replicated runs of our simulation model across 200 host generations. The top row displays time-series of host mean trait 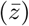, mean additive genetic value 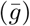, and mean additive microbial value 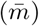. The bottom row displays time-series of host phenotypic variance (*P*), additive genetic variance (*G*), and additive microbial variance (*M*). The initial conditions are set to 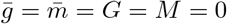, with completely even initial microbiomes, and simulation model parameters set to their defualt values described in Table 1.

Finite host population size also results in the chance associations between additive genetic and additive microbial values across host individuals. This happens for the same reason random genetic drift results in non-random associations of genetic alleles at different loci in the genome (i.e., linkage disequilibrium). This effect is most pronounced when microbial inheritance is entirely lineal (*𝓁* = 1), and when host population size is small. Hence, in our model, collective inheritance (corresponding to *κ >* 0) has analogous consequences as genetic recombination by leading to novel combinations of additive genetic and microbial values from standing variation. We therefore expect collective inheritance to buffer against the loss of diversity caused by host drift, and result in more accurate predictions of host trait dynamics by our analytical model. In the following section, we describe our approach to comparing our analytical and simulation models.

**Table 1:** Default parameter values used across simulations.

| Parameter | Symbol | Default Value |
| --- | --- | --- |
| Lineal Inheritance | $\ell$ | 1.0 |
| Collective Inheritance | $\kappa$ | 0.0 |
| Host Population Size | $N_e$ | $10^3$ |
| Microbiome Richness | $S$ | $10^2$ |
| Microbiome Community Size | $J_e$ | $10^6$ |
| Iterations of Microbiome Dynamics | $L$ | 30 |
| Strength of Selection | $s$ | 0.01 |
| Per Capita Additive Microbial Effects | $\alpha_i$ | See main text |

### Comparing Analytical and Simulation Models

To measure our analytical model’s ability to predict simulated host traits, we ran simulations for 50 host generations. At each host generation, we parameterized our analytical model using values from the simulation. We then compared the analytical model’s prediction to the simulated host mean trait value in the subsequent host generation by calculating an effect size based on Cohen’s *d* statistic (Cohen 1969; Sawilowsky 2009). In particular, writing 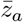 as the host mean trait predicted by our analytical model, 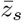 as the simulated host mean trait, and *P* as host trait variance, we compute the effect size

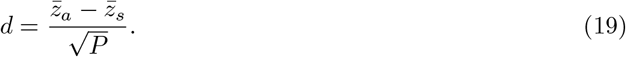

Because this effect size is normalized, we use it to make comparisons of our analytical model’s performance across different combinations of randomly drawn parameter values. We focus our comparisons on different modes of transmission, host population size, and number of microbe species. In Table 1, we list the default parameter values used across comparisons.

To simplify comparisons, we assume host trait variation is entirely explained by variation of additive microbial values such that *P* = *M* and *G* = *E* = 0. We assume selection on host traits is directional, with bigger trait values conferring greater fitness. In particular, we assume host fitness is given by *W* (*z*) = exp(*sz*), where *s >* 0 is the strength of selection. When the trait is normally distributed, as assumed by our analytical model, the selection gradient is given by *β* = *s*.

As summarized in Table 1, we use the following default values: lineal inheritance, *𝓁* = 1; collective inheritence, *κ* = 0; host population size, *N*_*e*_ = 1000; and microbiome richness, *S* = 100. For a given microbiome richness, *S*, we set *J*_*e*_ = 10^6^ and assume all microbiomes are initialized as completely even communities (where *n*_*i*_ = *J*_*e*_*/S* for each *i* = 1, …, *S*). We set the number of iterations of microbiome dynamics per host generation to *L* = 30. The strength of selection is fixed to *s* = 0.01. For each run of the simulation, we draw the per capita additive microbial effects *α*_1_, …, *α*_*S*_ independently and identically from a normal distribution with zero mean and variance given by

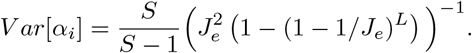

Using this distribution, the variance of microbiome dynamics (given by Equation (15)) will equal unity on average, given a completely even initial microbiome, making it comparable with the variance of mutation.

We applied this general setup to four different cases of randomly drawn parameters, with each case consisting of 1000 trials. In the first scenario, we drew host population size *N*_*e*_ at random such that *log*_10_(*N*_*e*_) ∼ Uniform(1,3). In the second, we drew microbiome richness *S* at random such that *log*_10_(*S*) ∼ Uniform(1,3). In the third, we drew collective inheritance *κ* at random such that *κ* ∼ Uniform(0,1), and set *𝓁* = 1 − *κ*. In the fourth case, we drew the total degree of microbial inheritance *κ* + *𝓁* at random such that *κ* + *𝓁* ∼ Uniform(0,1). To do so, we drew two independent variables, *X, Y*, uniformly distributed on the unit interval, and set *κ* = *XY* and *𝓁* = *X*(1 − *Y*).

## Results

In the section titled *Comparing Analytical and Simulation Models* above, we described our approach to measuring our analytical model’s ability to predict our simulation model. Results from this approach are given in Figure 4. Smaller effect size magnitudes are associated with better fits between our analytical and simulation models. The magnitude of the effect sizes can be interpreted using Sawilowsky’s heuristics (Sawilowsky 2009), and are summarized in Table 2. We find the effect sizes are symmetrically distributed around zero in each case, which shows that our analytical model’s predictions are not biased. The top left panel of Figure 4 shows that performance is worst when host population size is small, but rapidly increases with host population size. The top right panel of Figure 4 shows the analtyical model’s ability to predict the simulation model does not depend on microbiome richness. The bottom left panel shows performance decreases with levels of collective inheritance, but remains small according to Sawilowsky’s heuristics. In the bottom right panel, we find performance does not depend on the total level of microbial inheritance.

**Table 2:** Sawilowsky’s heuristics to interpret |*d*|. Smaller values are associated with agreement between analytical and simulation models.

| Value | Interpretation |
| --- | --- |
| 0.01 | Very Small |
| 0.2 | Small |
| 0.5 | Medium |
| 0.8 | Large |
| 1.2 | Very Large |
| 2.0 | Huge |

**Figure 4:**
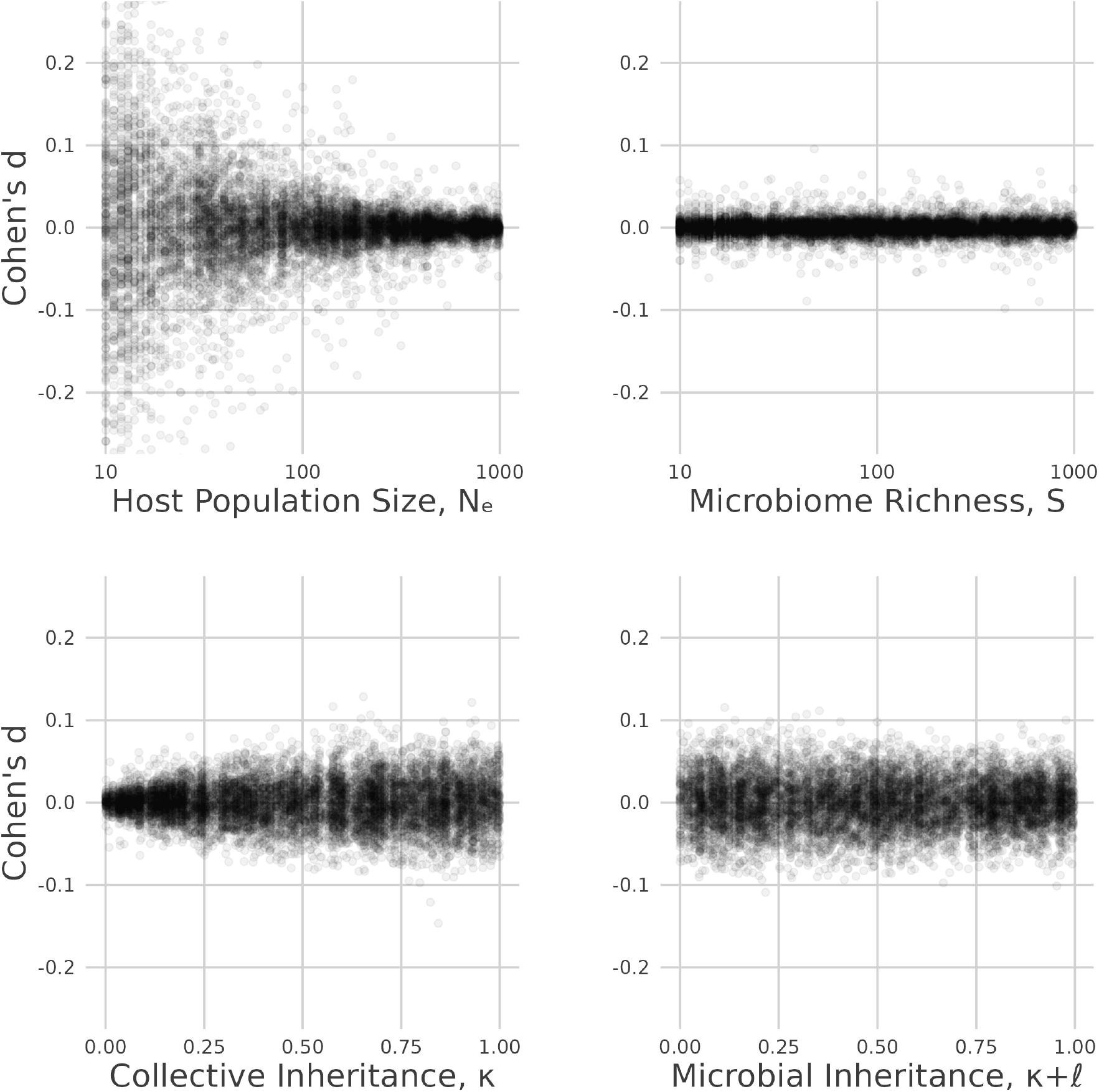
Performance of analytical model in predicting simulation model in four cases of randomly drawn model parameters. Each dot represents a comparison between the analytical model and simulation model over a single host generation. Positive values occur when the analytical model overestimates the simulated host mean trait. Negative values occur when the analytical model underestimates the simulated host mean trait.

## Discussion

Here, we introduced a simple mathematical model (Equation (10)) that provides analytical predictions of microbiome-mediated trait dynamics. Using a series of simplifying assumptions, we obtain an equation for the change of host mean trait in response to selection, lineal inheritance of genes and microbes, and environmental acquisition of microbes. By incorporating microbiome inheritance into the dynamics of host traits, our model formally generalizes a classical quantitative genetic expression for mean trait evolution (Lande 1979), and provides several novel insights into microbiome-mediated traits.

To test the robustness of our analytical model, we compared its predictions with microbiome-explicit simulations across different modes of transmission, number of microbe species, and host population size. Overall, we found our analytical model closely matched simulated host traits over a single generation when host population size is sufficiently large (≥ 100). We did not find any effect of microbiome species richness or total microbial heritability (*κ* + *𝓁*) on the performance of our analytical model. However, we found that increased collective inheritance slightly decreased our analytical model’s ability to predict simulations, but effect sizes remained small. This initial inquiry into our analytical model’s performance suggests that it is robust when critical assumptions are broken.

Our model also clarifies notions of heritability of microbiome-mediated traits. For instance, motivated by the definition of narrow-sense genetic heritability as the proportion of phenotypic variance explained by additive genetic effects, *h*^2^ = *G/P*, it is tempting to define a microbiome analog of *h*^2^ as the proportion explained by additive microbial effects, *M/P* . However, unlike additive genetic effects, which are assumed to comprise a subset of heritable genetic effects, additive microbial effects are not necessarily heritable. Under our model, inheritance of *M* is eroded when environmentally acquired microbes descend from lineages other than those found in the pooled parental microbiome. Hence, under our model, a more accurate definition for the narrow-sense microbial heritability of a microbiome-mediated trait is *b*^2^ = (*κ* + *𝓁*)*M/P*, the proportion explained by *heritable* additive microbial effects. Hence, the analogy between narrow-sense genetic heritability and narrow-sense microbial heritability only makes sense in the special case where all additive microbial effects are heritable by some combination of lineal and collective mechanisms, so that *𝓁* + *κ* = 1. However, unlike the interpretation of *h* as a path coefficient (Wright 1921; Lynch and Walsh 1998), whether or not *b* can be interpreted as a path coefficient is not obvious.

Our model also demonstrates the possibility of a host trait exhibiting a perfect response to selection (where offspring mean trait equals post-selected parental mean trait) in the absence of parent-offspring trait correlations (i.e., *Cov*(*z*^*′*^, *z*) = 0). This occurs in our model when additive genetic variance and environmental noise are absent, and when all microbes are collectively inherited, corresponding to *G* = *E* = 0 and *κ* = 1. Although such an extreme case is not likely to occur in nature, this scenario demonstrates that parent-offspring correlations underestimate the total heritability of microbiome-mediated traits, and highlights the adaptive potential of collective inheritance.

Although our model is sufficiently general to encompass classical quantitative genetics and microbiome inheritance, it has several important limitations. Firstly, we assume environmental acquisition of microbes draws from a shared, well-mixed pool. In reality, individuals are exposed to different environments, as reflected by the concept of an individual’s “exposome” (Wild 2012). Exposome variation can be modeled by replacing our single environmental pool of microbes with a number of distinct compartments, each with their own additive microbial value, that host individuals differentially sample. Correlated exposomes, where individuals are more likely to sample microbes from shared compartments, may be mediated by host attributes such as kinship and behavior. Another limitation of our model is its focus on a single host population. Naturally occurring host populations are spatially distributed. Replicating our model across several locations connected by host dispersal can provide insight into local adaptation of host-associated microbiomes and host-mediated spatial variation of environmental microbiomes (Meyer et al. 2018; Miller et al. 2018; Miller and Bohannan 2019). Our model also assumes that within-host microbiome dynamics happen independently across host individuals during development. A more realistic assumption would allow the exchange of microbes between host individuals directly via social contact and between host individuals and their environment continuously throughout host development. Finally, our model ignores interactions between host genome and microbiome in the determination of host phenotype. We discuss two distinct forms of these *G × M* interactions, and their similarity to related concepts, in the following paragraphs.

One form of *G × M* interaction involves host genetics mediating the phenotypic response to microbiomes. Treating host-associated microbiomes as components of the environment, this form of *G × M* interaction can be accounted for by combining an evolutionary reaction norm approach with a model of microbiome transmission. For instance, this can be done in our framework by implementing the host trait architecture *z* = *g*_1_ + *g*_2_*m* + *e*, where additive genetic values *g*_1_ and *g*_2_ are the intercept and slope of a linear reaction norm, and the additive microbial value *m* is an environmental variable. The dynamics of 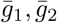, and 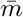 can then be obtained using a multivariate approach similar to that used by Lande (2009). Our model is obtained as the special case where reaction norm slope has no heritable genetic variation.

Although the *G × M* interaction described in the preceding paragraph illustrates similarities between microbiome-mediated traits and the evolution of reaction norms, there are two ostensible distinctions. Firstly, evolutionary reaction norm models typically consider phenotypic responses to a single macroenvironmental variable, such as regional temperature or precipitation, that is experienced by all individuals in a population or subpopulation (Via and Lande 1985; Gomulkiewicz and Kirkpatrick 1992; Gavrilets and Scheiner 1993; Chevin et al. 2010; Lande 2014). In contrast, additive microbial values comprise a set of microenvironmental variables unique to each host individual. Secondly, the environmental variables typically considered in reaction norm models are not transmitted between individuals, and thus have no heritable component. However, because the reaction norm concept can, at least in principle, be applied to traits responding to heritable microenvironmental variation, these distinctions are more a consequence of convention. In particular, microbiome-mediated traits can be regarded as a form of phenotypic plasticity in response to potentially heritable environmental variables.

Another form of *G × M* interactions involves host genetics mediating microbiome composition. This interaction can occur in the colonization of hosts by microbes, and in the dynamics of microbiomes during host development. Both of these modes can be incorporated into our framework by generalizing our analytical model. Microbial species sorting during host colonization can be incorporated by adding a selection component to the assembly process (with parameters considered as quantitative host traits in their own right) that biases the initial host microbial value *m*_0_. Host mediated microbiome dynamics can be incorporated by generalizing our model of microbiome dynamics. For example, unbiased microbiome dynamics, as assumed by our analytical model, can be obtained by modeling the within-host dynamics of additive microbial value as Brownian motion (Harmon 2019). A natural generalization is then to add a tendency of additive microbial value towards a value determined by host genotype, which can be done by replacing Brownian motion with an Ornstein-Uhlenbeck process (Vatiwutipong and Phewchean 2019), with parameters modeled as quantitative host traits.

Viewing host microbiome as an environmental factor, the *G × M* interaction described in the preceding paragraph is an example of niche construction (Odling-Smee et al. 2013; Scott-Phillips et al. 2014). The definition of niche construction requires the inheritance of natural selection pressures, called “ecological inheritance” (Odling-Smee et al. 2013). This occurs in our model when focusing on selection of host genotype. The transmission of microbiomes across host generations alters host trait architecture which, consequently, alters selection on host genotype, forming a mode of ecological inheritance. Hence, our model of microbiome-mediated trait dynamics can be applied as a quantitative genetic approach to study a particular form of niche construction that complements previous approaches (Fogarty and Wade 2022).

## Conclusion

In this paper, we introduced a novel approach to model the response of microbiome-mediated traits to selection, and obtained a generalization of a classical quantitative genetic expression that accounts for microbiome transmission. By comparing with microbiome-explicit simulations, we found that our analytical predictions are robust to finite host population size, species depauperate microbiomes, and variable modes of microbiome transmission. Leveraging insights from our analytical model, we suggested a definition for the narrow-sense microbial heritability of a microbiome-mediated trait, and discussed connections with phenotypic plasticity and niche construction. Taken together, this work offers first steps towards establishing a theory of microbiome-mediated trait dynamics, and highlights the adaptive potential of host-associated microbiomes.

